# SPDE: A Multi-functional Software for Sequence Processing and Data Extraction

**DOI:** 10.1101/2020.11.08.373720

**Authors:** Dong Xu, Zhuchou Lu, Kangming Jin, Wenmin Qiu, Guirong Qiao, Xiaojiao Han, Renying Zhuo

## Abstract

Efficiently extracting information from biological big data can be a huge challenge for people (especially those who lack programming skills). We developed **S**equence **P**rocessing and **D**ata **E**xtraction (SPDE) as an integrated tool for sequence processing and data extraction for gene family and omics analyses. Currently, SPDE has seven modules comprising 100 basic functions that range from single gene processing (e.g., translation, reverse complement, and primer design) to genome information extraction. All SPDE functions can be used without the need for programming or command lines. The SPDE interface has enough prompt information to help users run SPDE without barriers. In addition to its own functions, SPDE also incorporates the publicly available analyses tools (such as, NCBI-blast, HMMER, Primer3 and SAMtools), thereby making SPDE a comprehensive bioinformatics platform for big biological data analysis.

**Availability:** SPDE was built using Python and can be run on 32-bit, 64-bit Windows and macOS systems. It is an open-source software that can be downloaded from https://github.com/simon19891216/SPDEv1.2.git.

## Introduction

Extracting and organizing valid information from biological big data are the two foundations of bioinformatics analyses. In recent years, the rapid accumulation of genome data^1,2^ indicates that more efficient and reliable tools are needed to extract and organize the resultant biological big data. Although a lot of bioinformatic software^3,4^ have been developed, many are distributed in Linux systems or require users to have a knowledge of programming language. This poses huge challenges, especially for Windows users, and limits the ability of researchers to extract useful information from big data. Therefore, data extraction software is urgently needed to make up for this deficiency.

Gene family and omics analyses are two import aspects in bioinformatics analysis^5,6^. Extracting sequences (e.g., full-length gene sequences, promoter sequences and untranslated regions sequences (UTR)) from big data is the basis of gene family analysis^7^. The coding sequences (CDS) and protein sequences can be obtained easily (as the partial results of the genome sequencing), for example from Phytozome, the Plant Genomics Resource (https://phytozome.jgi.doe.gov/pz/portal.html). However, obtaining promoter sequences, full-length gene sequences and UTR can be difficult and time-consuming, especially when a gene family has more than 100 members. The extraction of effective information is the premise of omics analyses. For example, in transcriptome analysis, the main focus is often on genes that are differentially expressed under different experimental treatments^8^. It is also important to generate data in a file format that can be input to downstream software. For example, Circos^9^, which is a visualization tool for genomic information, has strict requirements for the format of input files (http://circos.ca/documentation/tutorials/configuration/data_files/). Although many Circos tutorials are publicly available, very few software can be used directly to generate files in the correct format.

The analysis software, NCBI blast+ (ftp://ftp.ncbi.nlm.nih.gov/blast/executables/blast+/LATEST/), ClustalW^10^, Muscle^11^, MAFFT^12^, HMMER (http://hmmer.org/), MCScanX^3^, and Primer3^13^, are important for sequence alignment, detecting homologous genes, collinearity analysis, and primer design; however, they can be used only through programming or command lines. Moreover, the input file of Primer3 (not the online version) needs to be in a specific format. Therefore, to utilize them skillfully and effectively can be a challenge for researchers, especially for wet-lab biologists.

To address the practical problems in sequence processing and data extraction, we developed **S**equence **P**rocessing and **D**ata **E**xtraction (SPDE) as an integrated tool for sequence processing and data extraction. The main goal was to minimize the amount of time required to deal with big data files. Thus, except for the first module, the other six modules in SPDE use batch processing. For example, except for the required files, users need to provide only the gene identifiers (ids) to extract the full-length gene sequences and their promoter sequences from genomic data. In addition, we simulated the awk, sed, and other Linux commands to develop a new extraction engine. To extract information from a big data file, users need to provide only rows/columns ids or other filter criteria, then the module will write required the rows or columns into a new file. The interfaces of NCBI blast+, ClustalW, Muscle, MAFFT, HMMER, MCScanX, and Primer3 also have been integrated in SPDE. The aim was to provide a tool that allows researchers to focus on the meaning of the bioinformatics data rather than on how to obtain relevant information.

## Methods and Features

### Overview of SPDE functions

The modules and some functions of SPDE are shown in Fig. 1. Currently, SPDE contains nine modules comprising 100 functions that range from processing a single gene sequence to extracting genomic information.

**Fig. 1.**
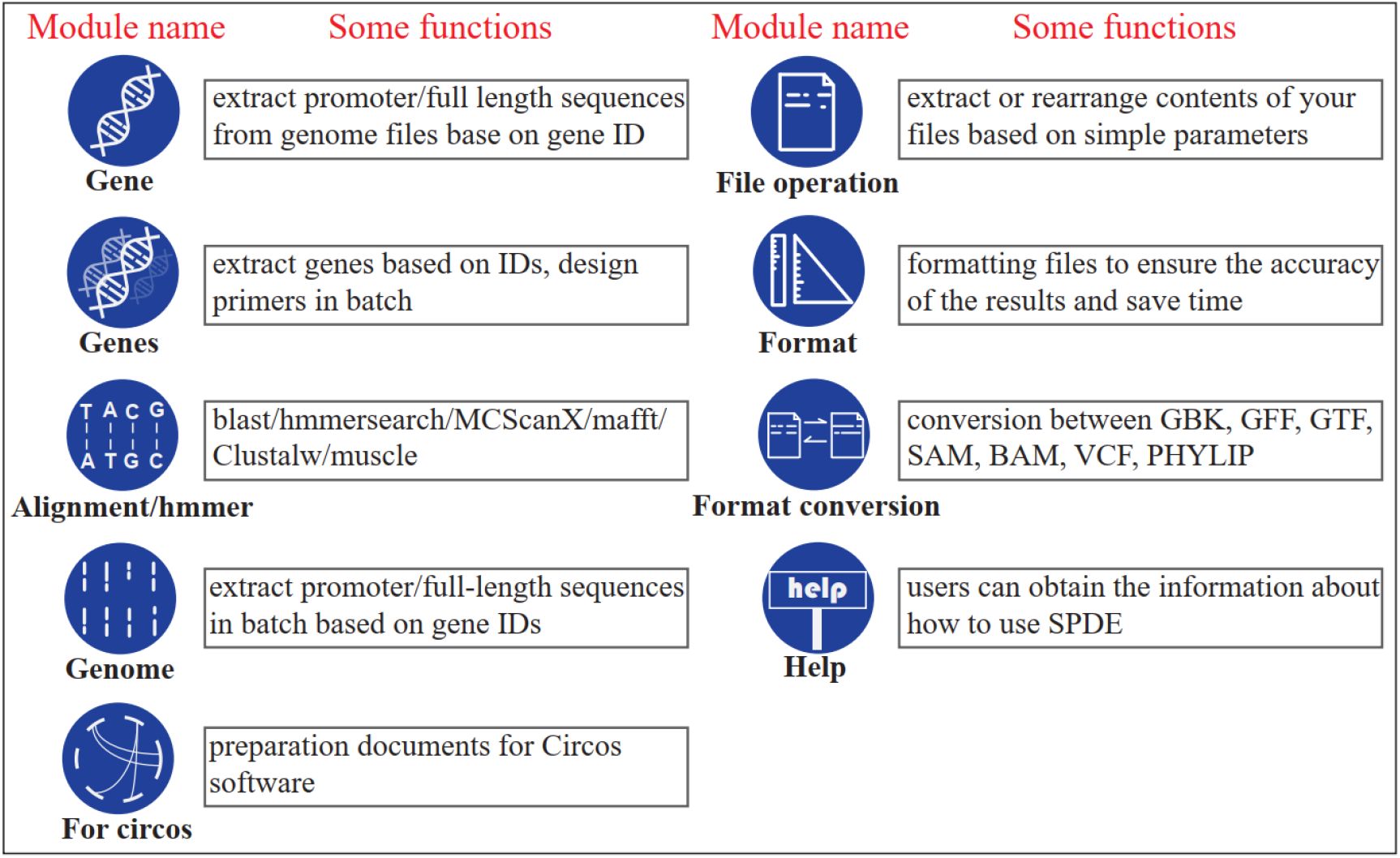
Overview of SPDE functions. Functions of nine modules in SPDE are ranging from sequence mining to organizing genome information, which mainly play roles in gene family analysis and genome analysis.

The functions of ‘Gene’ module can help biologists mine promoter sequences, full-length sequences and UTR sequences from genomic files, based on gene ID. Then, ‘design primers’ can be used for generating primers. Meanwhile, sequence processing (such as, reverse complement and translation) was also set in this module (Supplementary video 1). In the ‘Genes’ module, SPDE performs batch processing of multi-gene files, such as batch primer design and translation (Supplementary video 2). In the ‘Alignment/HMMER’ module, we developed a user-friendly interface for Blast, HMMERsearch, ClustalW, Muscle, MAFFT (for Windows users) and MCScanX. The 13 functions in this module can be used to complete alignment, collinearity analysis, and homologous genes identification (Supplementary videos 3.1 and 3.2). The ‘Genome’ module can be used to compile statistics of genome information, such as counting the numbers and positions of genes on each chromosome, and to extract promoter and full-length gene sequences in batches (Supplementary video 4). The ‘For circos’ module automatically generates files that are formatted for use in Circos software from input files in a different format (Supplementary videos 5.1 and 5.2). The ‘File operation’ module allow users to replace and delete the content of input files, and to extract, rearrange, and cut file contents, according to their requirements. All of these functions can be achieved by setting simple parameters (Supplementary videos 6.1 and 6.2). Although strict file formats have been specified for bioinformatics applications (such as fasta and General Feature Format (gff)), we found subtle differences in the format of files, even among files of the same format type. Unfortunately, these subtle differences seriously affected the processing efficiency and accuracy of SPDE. Therefore, we developed the ‘Formatting’ module so that users can format their files according to the prompts of the SPDE interface, before using SPDE for their analyses. The last module is ‘Format_conversion and others’. This module supports conversion between files in different formats (Supplementary videos 7).

### Information extraction

Three types of information can be extracted using SPDE: sequence information, genomic information, and rows/columns information. For sequence extraction, except for the required files, users need to provide only gene ids to obtain the CDS, UTR, promoter, and full-length gene sequences from CDS files (Supplementary video, Genes), genome-sequence files and gff/gff3 files (Supplementary videos, 1 and 4), respectively. Extracted genomic information includes chromosome/scaffold ids, length, GC content, starting and end positions of genes, and numbers of introns, exons, and genes (Supplementary video4). To focus on partial chromosomes/scaffolds instead of the whole genome, we provided related functions that allow users to extract information from a gff/gff3 file by inputting the chromosome/scaffold id (Fig. 2A). The output file can be used as the input file to extract the information of a specific chromosome/scaffold (Fig. 2B).

**Fig. 2.**
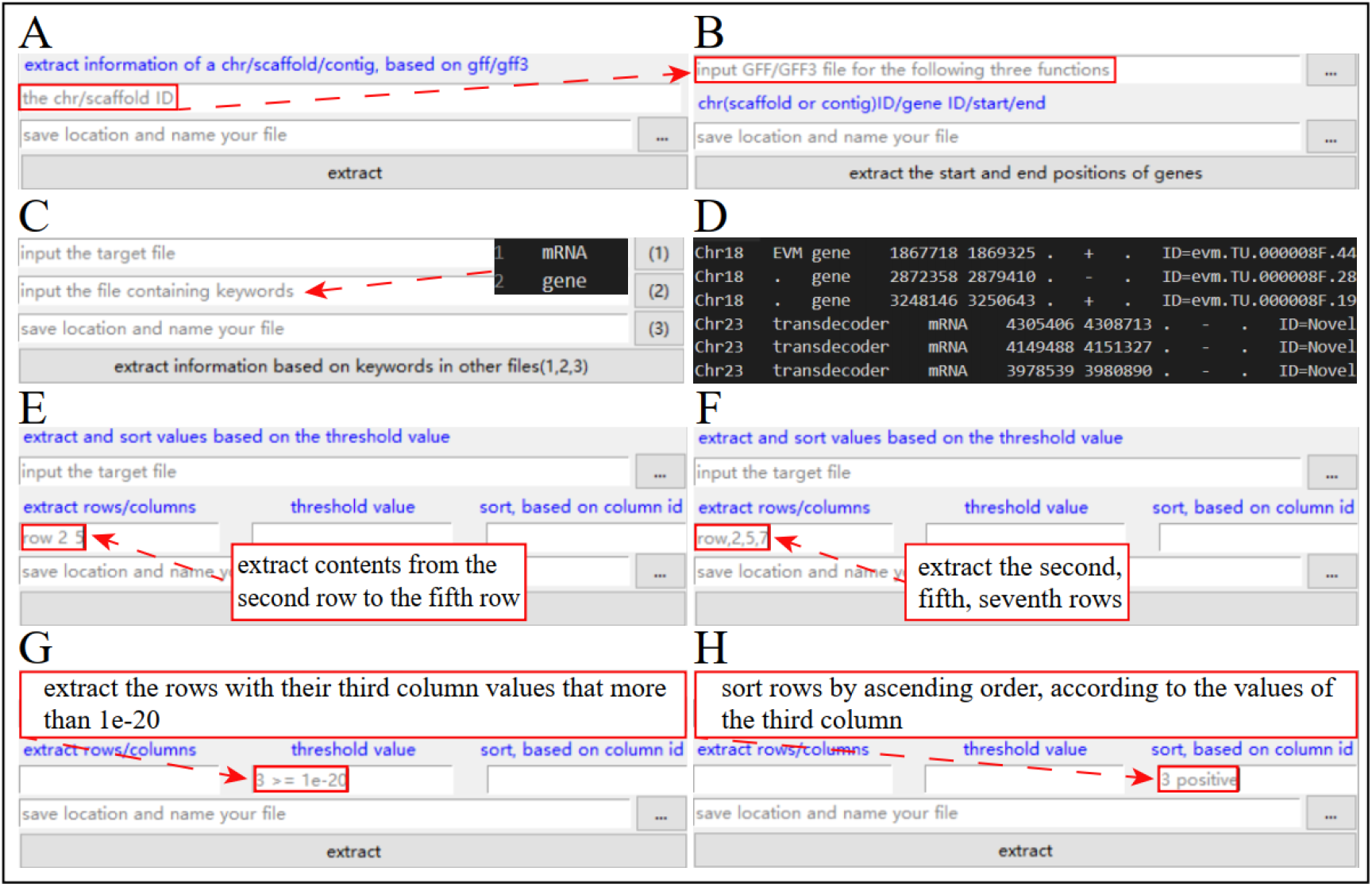
Information mining. (A) SPDE can extract partial chromosomes/scaffolds information from gff/gff3 files by inputting the chromosome/scaffold identifier (id). (B) The output file from (A) can be used as the input file to extract information about the genes that are located in the extracted chromosome/scaffold. (C, D) A file containing keywords can be set up to extract information from user files. (E, F) Setting parameters separated by a space or a comma will extract continuous or discontinuous rows/columns, respectively. (G) Setting a threshold value will extract eligible rows. (H) Setting ‘positive/negative’ will sort the content of the file obtained in (G) in ascending/descending order.

The rows/columns extraction module allows the same document to be interpreted from different perspectives because files will often contain a lot of information that is not used in subsequent analysis. For example, the focus may be on differentially expressed genes before and after a particular experimental treatment. Furthermore, opening big data files in the existing Office software of Windows can be difficult, let alone dealing with them, and big data files are becoming more and more common in omics. Using the rows/columns extraction module, according to their own needs, users can easily design parameters to extract information from their files instead of opening them (Supplementary videos 6.1 and 6.2). Moreover, we set the function of big file preview to help users determine the required ids (Fig. S1). There are nine functions in this module. For example, keywords can be set up (Fig. 2C) to extract the required content (Fig. 2D). In the input file, the only requirement is that the columns are separated by a Tab. There are eight ways to complete the extraction process. To extract continuous columns/rows, the parameters should be separated by a space (Fig. 2E), whereas, to extract discontinuous columns/rows, the parameters should be separated by a comma (Fig. 2F). To extract rows by setting threshold values, for example, to obtain differentially expressed genes with expression levels above or below a particular threshold cutoff, threshold values can be set as shown in Fig. 2G. To sort the content of a new file after extracting the required rows, the parameters can be set as shown in Fig. 2H.

### Creating input files suitable for downstream software

Our files sometimes may not be suitable for downstream software analysis because of (1) subtle differences in same-type files (Fig. S2A); (2) differences in format between the original files and input file of downstream software (Fig. S2B); and (3) extra information in original the files that is not required for the follow-up analysis (Fig. S2C). In SPDE, four functions were set up to solve such problems (Fig. 3). To delete specific strings, the function can be completed as shown in Fig. 3A (Supplementary video 6.1). To remove extra information from the original file, the functions can be set up as showed in Fig. 3B and C. To reorganize the file content to meet the format requirement of downstream software, the related parameters can be set up as shown in Fig. 3D (Supplementary videos 6.2).

**Fig. 3.**
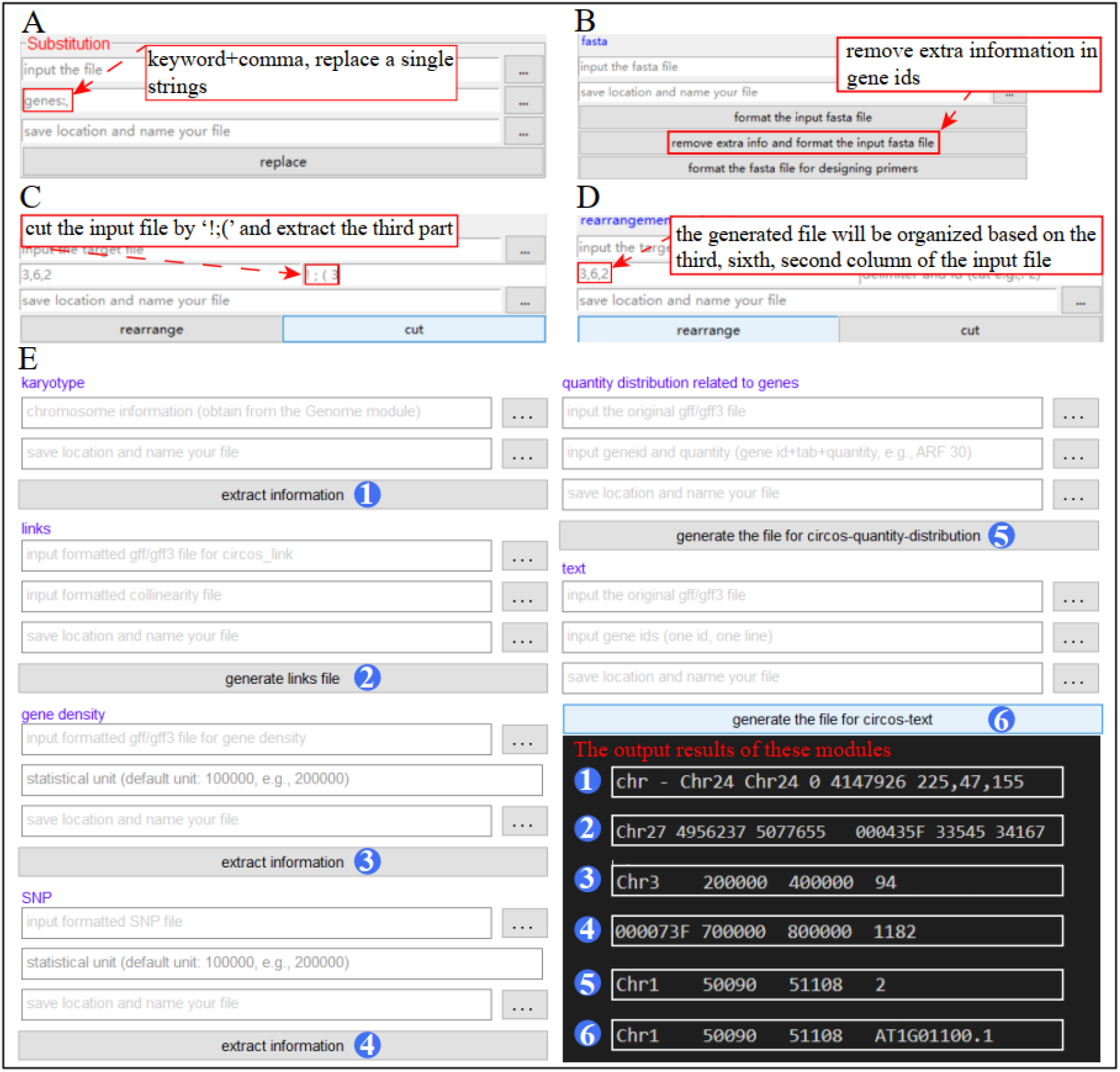
Creating input files for downstream software. (A) Function to remove specific strings from original files. (B, C) Function to remove the extra information from original files. (D) Function to reorganize user files to meet the format requirement of downstream software. (E) Functions for generating input files for the Circos software.

In particular, considering the importance of the Circos software in displaying genomic information, we created a ‘For circos’ module in SPDE to generate the output files which meet the format required for Circos input files (Supplementary videos 5.1 and 5.2). As shown in Fig. 3E, this module can produce six types of files (karyotype, links, gene density, heatmap, text, SNP) that can be used directly as input files for Circos.

### Data format conversion

Specific types and formats of the input data should meet the requires of different data extraction and machine learning algorithms^14^. Thus, format conversion is an operation that we often need to perform in the process of bioinformatics analysis. In SPDE, nine functions were set up for helping users convert their files to another format (Fig. 4). For example, the function of ‘convert fastq to fasta’ (Fig. 4A) can change the input file to a fasta file (Supplementary video 2). Two kinds of files (GenBank files and gff/gff3 files) can also be converted to each other (Fig. 4B, Supplementary video 6.1). With the help of SAMtools^15^, Sequence Alignment/Map format files (sam files) will be changed to Binarysequence Alignment/Map format files (bam files) or Variant Call Format files (VCF files, Fig. 4C). Furthermore, the VCF files can also be transformed to another format files^16^ (in Windows versions, Fig. 4D, Supplementary video 7).

**Fig. 4.**
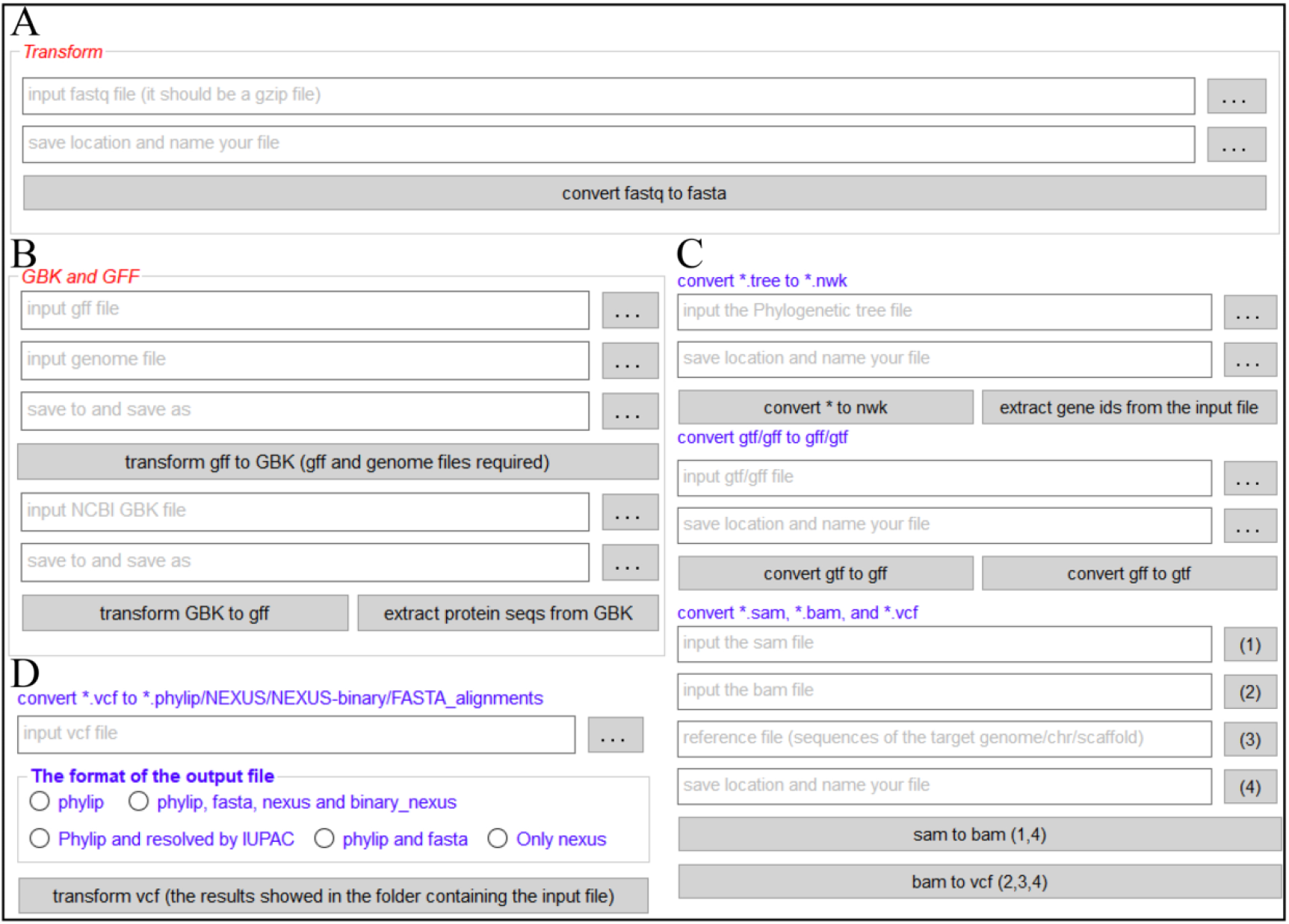
File format conversion. A, fastq to fasta; B, conversion between (GenBank files and gff/gff3 files); C, phylogenetic tree file to NEWICK file, conversion among sam files, bam files and VCF files; D, VCF files to another format file.

### Using the user-friendly SPDE interface

The user-friendly SPDE interface used point-and-click operations. Although the software NCBI blast+, HMMERsearch, ClustalW, Muscle, MCScanX, and Primer3 (Fig. 5) are excellent tools for sequence alignment, homologous genes identification, collinearity analysis, and primer design, the required command-line operations can make them difficult for users to implement. For this reason, we developed interfaces for these programs in SPDE whereby users can obtain the results by inputting their files according to the prompts of the interface (Supplementary videos 3.1 and 3.2).

**Fig. 5.**
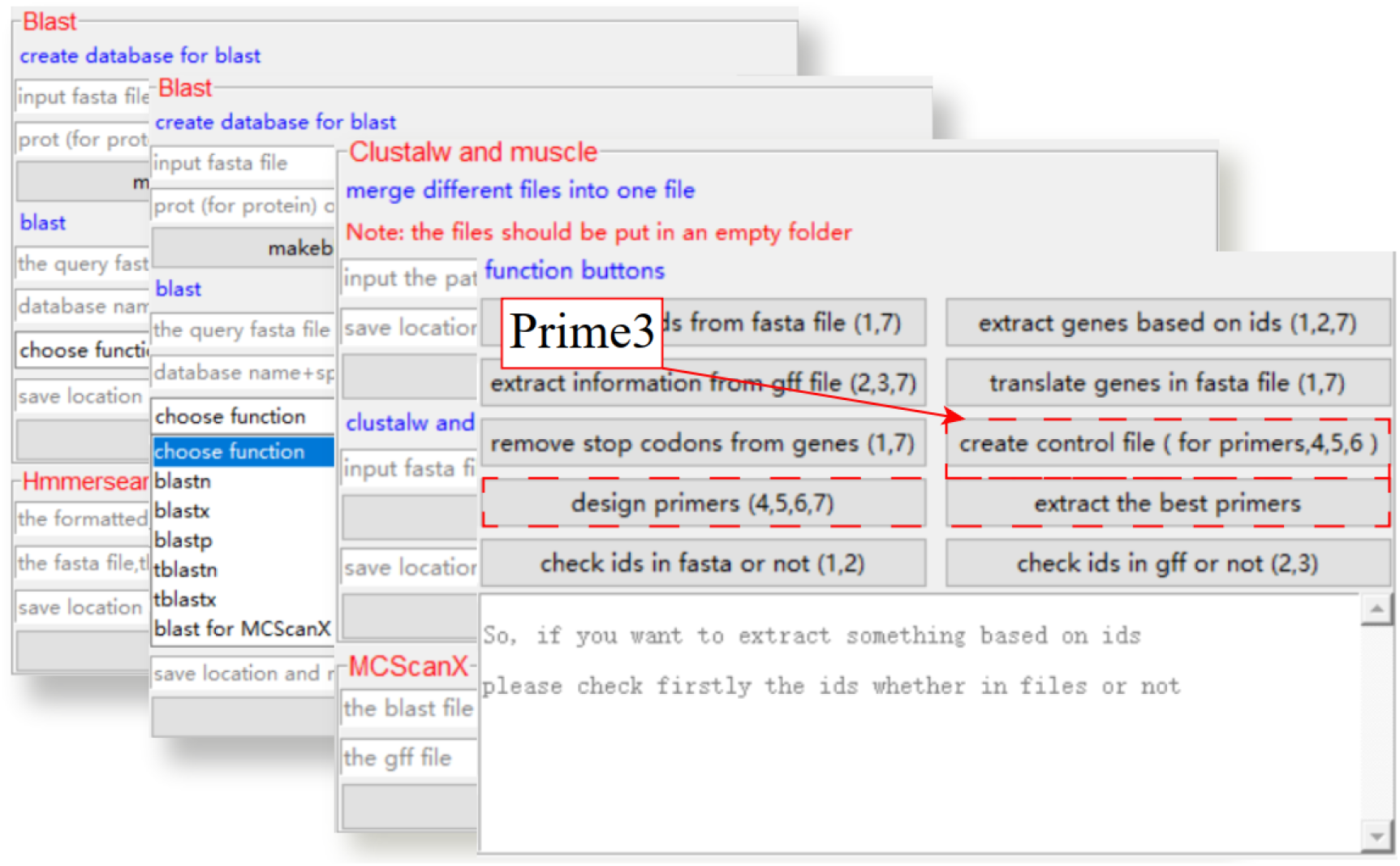
SPDE incorporates NCBI blast+, HMMERsearch, MCScanX, ClustalW, Muscle and Primer3. HMMERsearch, MCScanX, ClustalW, Muscle, NCBI blast+ and Primer3 can be used without the need for programming or command lines. All users need to do is entering the corresponding content as prompted and clicking the relative buttons.

Considering that the design of primers for multiply genes or sequences is a common task, we created the ‘Genes’ module to finish it (Supplementary video 2). The settings for the functions in this module are shown in Fig. 5. The ‘create control file (for primers,4,5,6)’ and ‘design primers (4,5,6,7)’ buttons are used to generate primers. A total of five pairs of possible primers is recommend. Then, the ‘extract the best primers’ button can be used to extract the best primers for a particular purpose (Fig. S3).

### Usability of SPDE

There is a ‘Help’ button or ‘About SPDE’ button at the SPDE interface that opens a list of module names that can be clicked to obtain the help document for the selected module, and article attachments that contain videos showing the use of SPDE also can be downloaded (Supplementary video 8). Although, the interface of different versions (32-bit, 64-bit Windows and macOS version) may have slight differences, there is enough prompt information on the software interface to help users apply SPDE. For example, the prompt ‘input formatted genome sequences’ tells users where the formatted genome sequences should be input (Fig. 6, Supplementary video 4). Users can use functions in the ‘Formatting’ module to format genome sequences that have not been formatted. Although formatting files may take some time because the operation objects are big data files, we found that ‘formatting files’ was an effective way to improve the efficiency of SPDE processing.

**Fig. 6.**
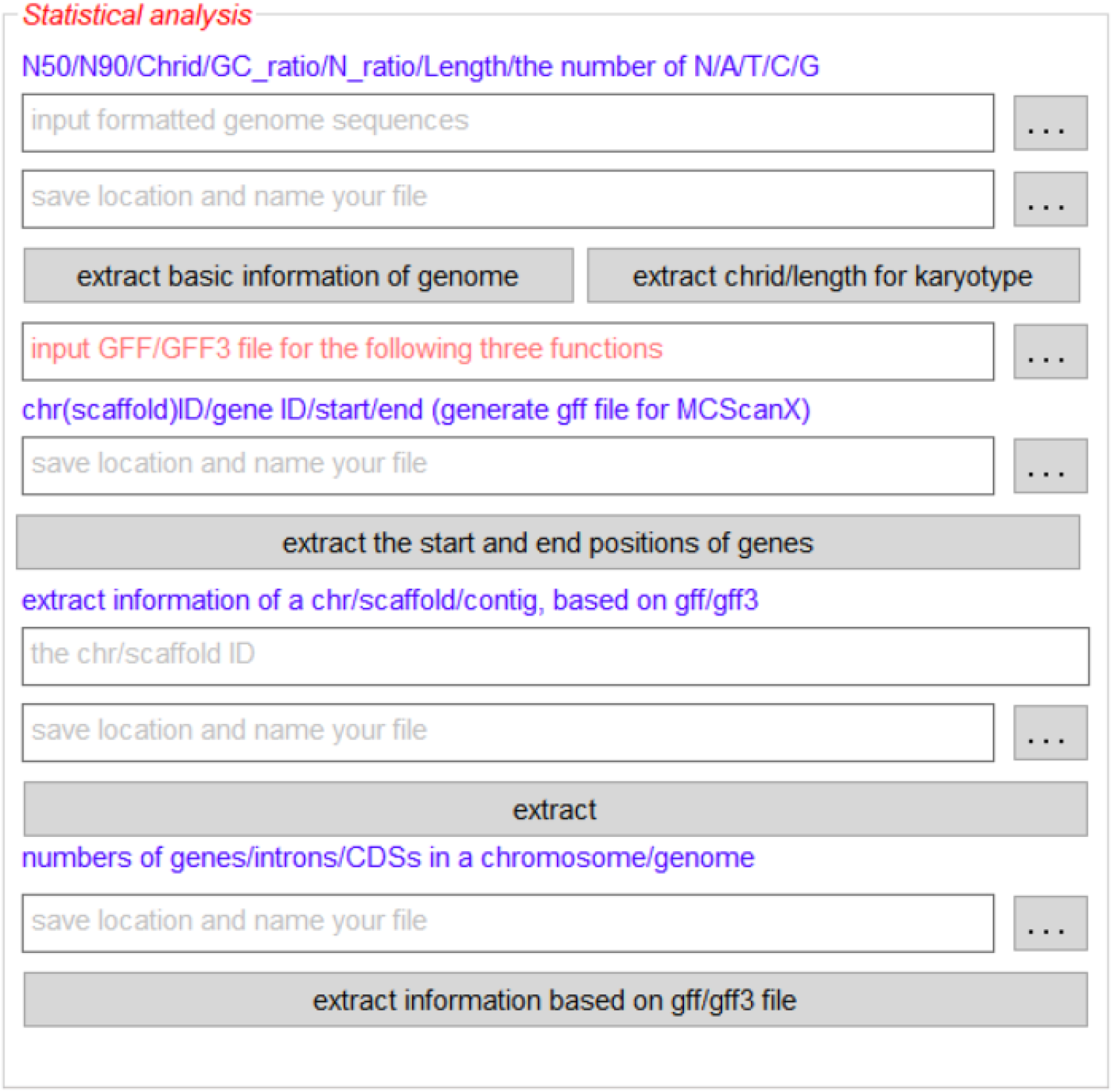
Enough prompt information on the software interface. To remove barriers in the use of SPDE, three kind of prompt information were left in the interface. The red character is the title or the warning message; the blue character is the function introduction or the contents of results; the grey character is the content that should be input in the box.

## Conclusions

SPDE has an important role in facilitating sequence processing and data extraction. As we gradually improve its functions, SPDE will become a powerful tool for biologists in processing biological big data.

## Acknowledgments

This work was supported by a grant from National Transformation Science and Technology Program (2018ZX08020002-005-003), the Zhejiang Science and Technology Major Program on Agricultural New Variety Breeding (No. 2016C02056-1), the National Natural Science Foundation of China (No.31870647 and NO.31872168).

